# Integrating Dynamic Network Analysis with AI for Enhanced Epitope Prediction in PD-L1:Affibody Interactions

**DOI:** 10.1101/2024.02.08.579577

**Authors:** Diego E.B. Gomes, Byeongseon Yang, Rosario Vanella, Michael A. Nash, Rafael C. Bernardi

**Author notes:** This authors contributed equally to this work (Diego E.B. Gomes, Byeongseon Yang).

## Abstract

Understanding binding epitopes involved in protein-protein interactions and accurately determining their structure is a long standing goal with broad applicability in industry and biomedicine. Although various experimental methods for binding epitope determination exist, these approaches are typically low throughput and cost intensive. Computational methods have potential to accelerate epitope predictions, however, recently developed artificial intelligence (AI)-based methods frequently fail to predict epitopes of synthetic binding domains with few natural homologs. Here we have developed an integrated method employing generalized-correlation-based dynamic network analysis on multiple molecular dynamics (MD) trajectories, initiated from AlphaFold2 Multimer structures, to unravel the structure and binding epitope of the therapeutic PD-L1:Affibody complex. Both AlphaFold2 and conventional molecular dynamics trajectory analysis alone each proved ineffectual in differentiating between two putative binding models referred to as parallel and perpendicular. However, our integrated approach based on dynamic network analysis showed that the perpendicular mode was significantly more stable. These predictions were validated using a suite of experimental epitope mapping protocols including cross linking mass spectrometry and next-generation sequencing-based deep mutational scanning. Our research highlights the potential of deploying dynamic network analysis to refine AI-based structure predictions for precise predictions of protein-protein interaction interfaces.

## Introduction

Immunotherapies are a potentially transformative class of cancer therapy, with checkpoint inhibition representing a leading strategy.^1^ The efficacy of such therapies depends on achieving precise molecular interactions between a therapeutic biomolecule and the correct epitope on the checkpoint protein. In the case of targeted PD-L1 therapies (Fig. 1A) with demonstrated efficacy against a range of cancers,^2–5^ anti-(PD-L1) antibodies must bind the PD-L1 extracellular domain (Fig. 1B, UniProt Q9NZQ7) in a manner that blocks native PD-1 ligand binding in order to effectively disrupt cancer cells’ evasion of the immune response. The development of novel therapeutics therefore relies on achieving precise and specific binding to a particular surface epitope. This highlights the urgent need for efficient methods to rapidly and accurately identify binding epitopes of therapeutic biomolecules on the surfaces of immune checkpoint proteins.

**Figure 1:**
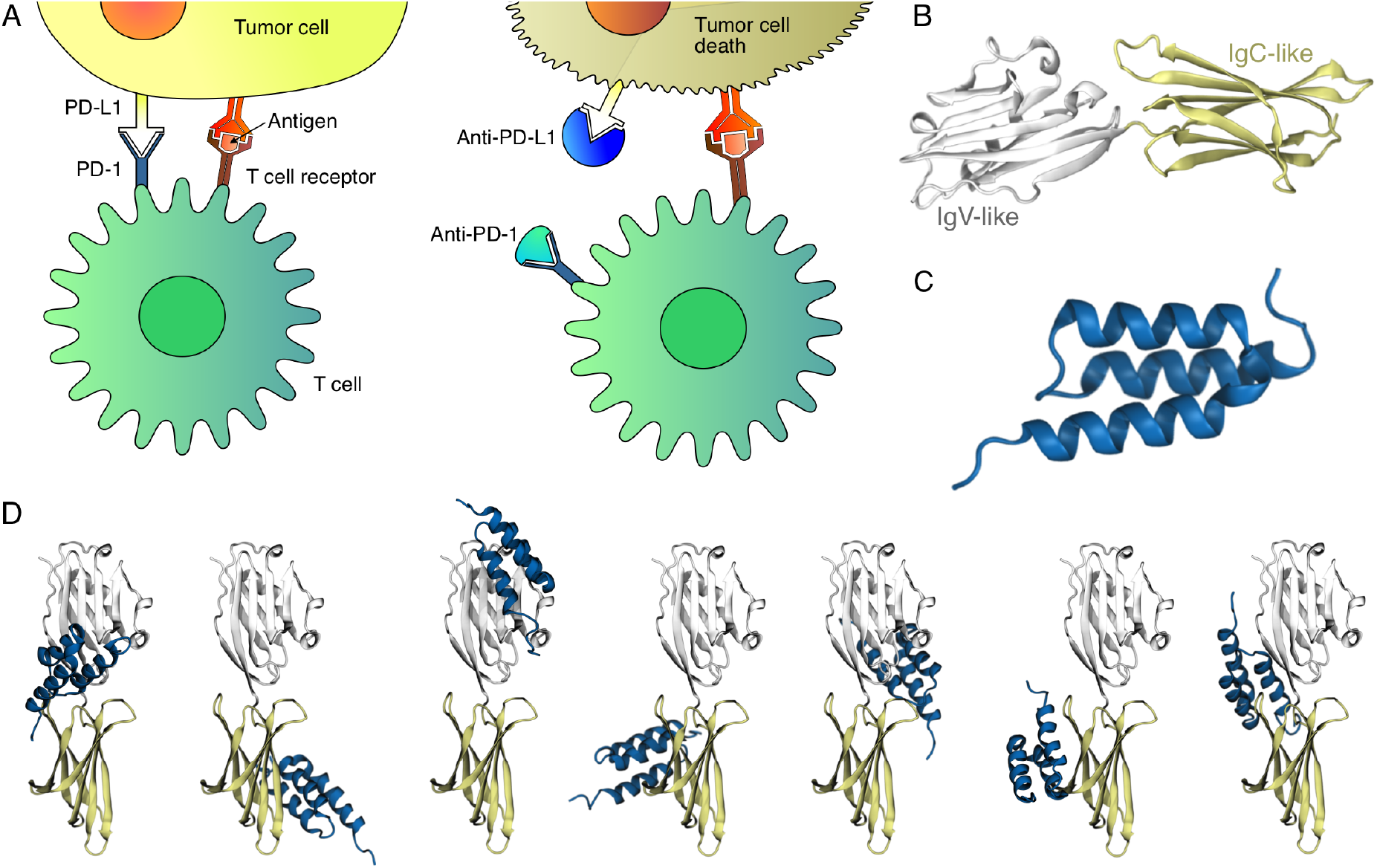
Overview of PD-L1 cancer targeting and lack of reliability of blind docking. A) (Left) By binding to PD-1, tumor cells overexpressing PD-L1 are able to inhibit T-cell activation, thereby avoiding clearance. (Right) Interfering with PD-L1 interactions using an immune checkpoint inhibitor (e.g., anti-(PD-L1) or anti-(PD-1) antibody) empowers T-cells to effectively eliminate tumor cells. B) Structure of PD-L1 (UniProt: Q9NZQ7) with its two domains: IgV-like (white) and IgC-like (yellow). C) Cartoon representation of the 3-helix bundle Affibody. D) Blind docking predictions for the complex using ClusPro 2.0 server ^34–38^ show a diverse set well ranked solutions.

Full length IgG monoclonal antibodies are a mainstay of cancer immunotherapy, but many IgG formulations suffer from limitations such as limited tissue penetration, suboptimal pharmacokinetics, and complex post-translational modifications that must be optimized during production. These challenges have spurred research into alternative protein scaffolds for PD-L1 targeted therapies, including diabody, DARPin, and affibody scaffolds.^6–8^ Such non-antibody scaffold proteins can be engineered using *in vitro* directed evolution to isolate high affinity binding domains against a range of molecular targets.^9–11^ With a molecular weight of ∼6.5 kDa, the highly stable affibody triple *α*-helix bundle ^12,13^ (Fig. 1C) in particular offers a promising alternative scaffold. For epitope mapping of antibodies and non-antibody scaffolds, a variety of experimental methods have been employed, including X-ray crystallography, NMR, mass spectrometry, peptide arrays and deep mutational scanning-based approaches.^14–22^ Computational methods have potential to be faster and less costly, however, even advanced artificial intelligence (AI)-based AlphaFold and RoseTTAFold ^23–26^ have faced challenges in accurately predicting protein-protein interactions.^27–30^ While success cases exist, such as the model of a PD-L1/CD80 complex,^31^ large screening studies have shown a low number of highly-accurate antibody-antigen complex prediction.^32,33^ Similarly, traditional computational methods often fail to accurately identify binding epitopes, sometimes producing ambiguous results that suggest binding at multiple locations (Fig. 1D).

Here we present a comprehensive methodology for predicting and analyzing the structure and interactions of a PD-L1:Affibody complex,^39,40^ focusing on epitope mapping. Our approach relies on recent advances in both computational biophysics methodologies and the power of the latest generation of supercomputers.^41^ We employ an integrated computational approach that leverages AlphaFold, ClusPro, ^38^ and Zdock ^42^ for complex structure prediction, refines these structures through molecular dynamics simulations performed with QwikMD^43^ and GPU-accelerated NAMD,^44^ and employs dynamic network analysis for further insights.^45^ Experimental validation of our computational models is conducted using a suite of techniques including site-specific mutagenesis, biochemical assays, chemical cross-linking mass spectrometry, and next-generation sequencing (NGS)-based deep mutational scanning (DMS).^46^ Collectively, these methods confirm the computationally predicted protein-protein interaction interface and identify specific residues involved in the interactions.

## Results

We began by investigating the native PD-1:PD-L1 interaction interface, localized within the IgV-like domain of PD-L1. ^47^ An exhaustive analysis of bound PD-L1 structures in the Protein Data Bank (PDB) revealed insights into its interactions with other proteins and structural nuances of the complex assembly. By analyzing available PDB structures of PD-L1 bound to other proteins and focusing on amino acid residues within a 4Å radius of the binding interfaces, we found diverse sets of native contacts across PD-L1’s surface (Fig. 2A). This analysis highlights the complexity in defining the Affibody binding epitope on PD-L1 (Fig. 1D), and shows PD-L1’s promiscuity and the challenges faced by traditional blind-docking methods.

**Figure 2:**
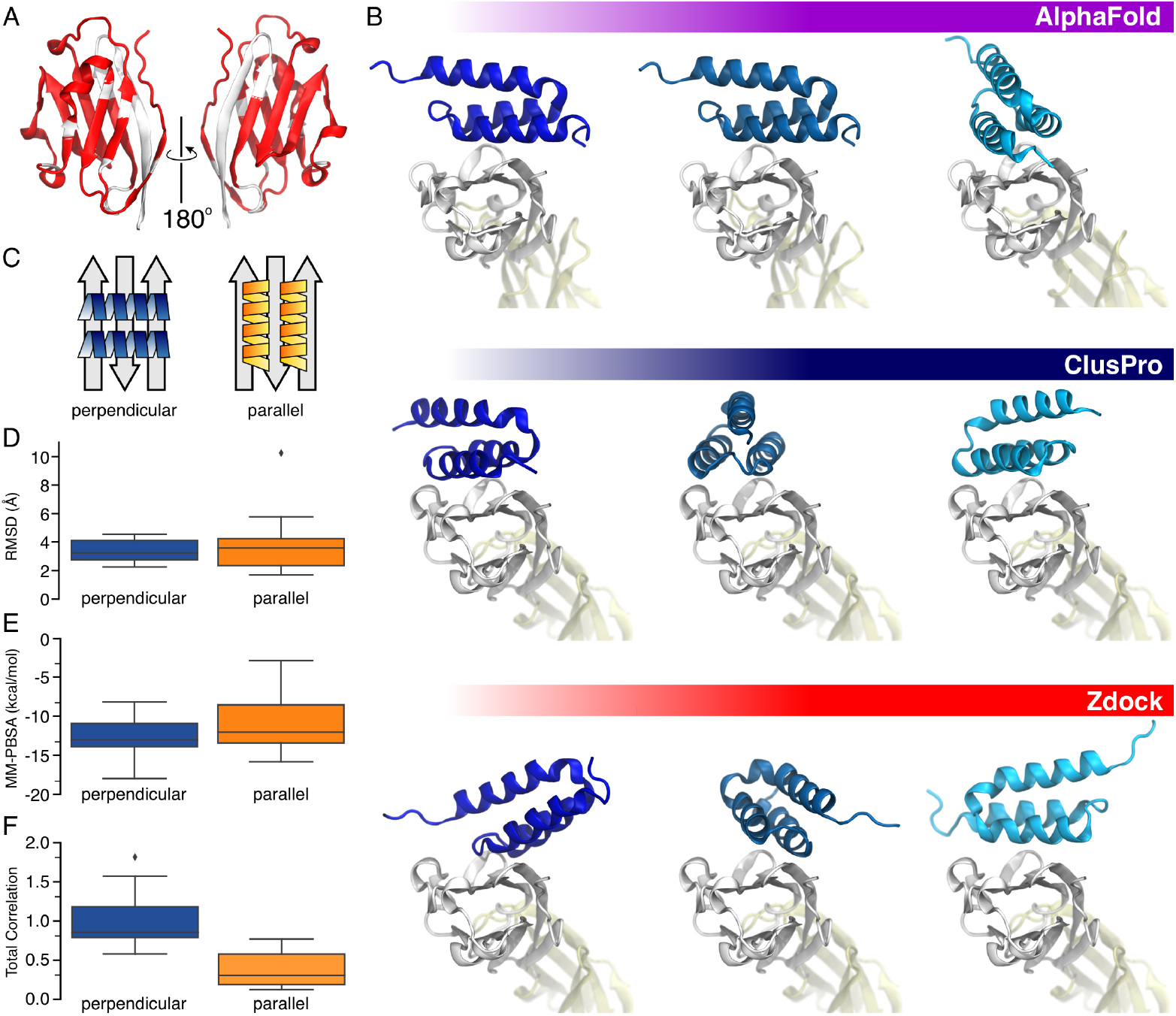
Results of structural predictions of the PD-L1:Affibody complex. A) Native contacts of PD-L1 complexes found in the PDB highlighted in red B) Comparative Analysis of PD-L1 and Affibody Binding Orientations: Alphafold2 Multimer, ClusPro, and ZDock Predictions. The top 3 ranked structures from each prediction method depict the binding orientations of PD-L1 and Affibody. Two dominant orientations were consistent across all predictions: one parallel to the beta sheet and another perpendicular to it. This highlights the convergence of computational approaches in capturing putative binding modes of this complex. C) Illustration of the two proposed orientations. D-F) Boxplots for independent properties obtained from sixteen 100-nanosecond all atom MD simulation replicates of the parallel and perpendicular orientations. D) Affibody RMSD after fitting to PD-L1. E) Estimation of binding free energy using the Molecular Mechanics Poisson-Boltzmann Surface Area (MM-PBSA) method. F) Sum of the correlation of interfacing residues normalized by the perpendicular average. The combined analysis of these MD-based metrics strongly supports the perpendicular binding mode between PD-L1 and Affibody as the most probable configuration, underscoring the robustness and consensus of computational assessment methods.

**Figure 3:**
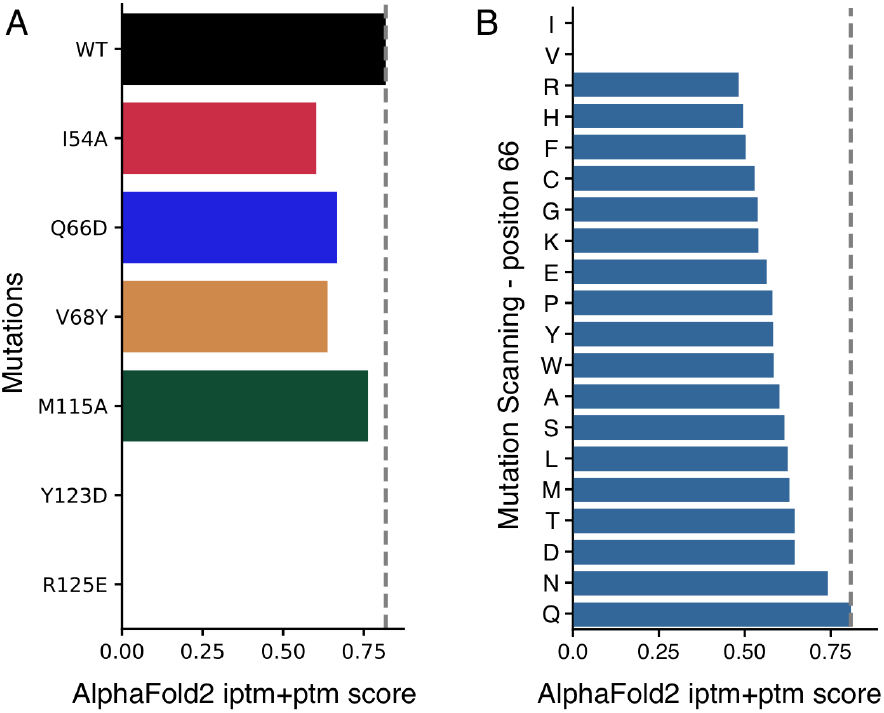
*In silico* site-specific mutagenesis to find PD-L1 residues involved in each of the binding modes. A) and B) AlphaFold2’s iptm+ptm score for the best ranking model of the complex. A) Screening of selected PD-L1 mutations within the predicted binding interface: I54A, Q66, V68Y, M115A, Y123D, and R125E. B) Computational mutational scanning for PD-L1 at position 66 for the perpendicular mode.

A potential alternative to traditional blind-docking approaches is provided by AI-based modeling tools, such as AlphaFold2, specifically its Multimer version.^23,24^ In this study, we carried out a comparative analysis of PD-L1:Affibody binding orientations using three distinct prediction methods: AlphaFold2 Multimer, ClusPro, and ZDock, with ClusPro and ZDock utilizing existing structural data for binding space constraints and AlphaFold2 Multimer operating in a “blind-mode” (Fig.2B). The top three predicted structures from each method revealed two main binding orientations for the Affibody: a parallel and a perpendicular orientation of Affibody relative to PD-L1’s beta sheets (Fig.2C). Although dual binding modes have been reported for certain receptor:ligand complexes, ^48,49^ it is rather an exceptional case, and the parallel and perpendicular binding orientations for PD-L1:Affibody complexes were considered ostensible. The output from the three static computational methods could therefore not confidently determine the binding epitope on PD-L1, and we sought to further analyze the structures to determine which orientation is more probable.

To investigate the stability of the two potential PD-L1:Affibody binding orientations, we next conducted molecular dynamics (MD) simulations using GPU-resident NAMD 3.0, ^44^ running 16 independent 100 *ns* replicas for each orientation. Initial visual assessments using VMD^50^ indicated greater stability in the perpendicular orientation (Fig. S1). For a more rigorous evaluation, we analyzed the final 25 *ns* of each trajectory using three methods. Firstly, we measured the root-mean-square deviation (RMSD) of the Affibody from its initial conformation, finding similar mean RMSD values for both orientations but with larger fluctuations in the parallel mode (Fig.2D). Secondly, the Molecular Mechanics Poisson-Boltzmann Surface Area (MM-PBSA) method was used to calculate binding free energy, yielding comparable results but greater variability for the parallel orientation (Fig.2E). Lastly, leveraging a technique developed by our group we employed generalized-correlation-based Dynamic Network Analysis ^45^ to derive relative binding strengths.^51^ The results, displayed in Fig.2F and Fig.S2, reveal the perpendicular mode as notably more stable. These combined findings suggest the perpendicular binding conformation as the most probable for the PD-L1:Affibody interaction.

### Experimental Validation of Predicted Epitope

To validate the predicted binding mode, we first used VMD^50^ for visual inspection and AlphaFold2 Multimer scores to identify mutations to PD-L1 that would potentially disrupt Affibody binding. Mutations like I54A and Q66D significantly reduced the AlphaFold2 structure score, suggesting impaired binding (Fig.A, Fig.S3). A detailed analysis at position 66 showed that the original amino acid Glu (Q) favored binding, while positively charged amino acids such as Arg (R) and His (H) were detrimental (Fig.B, Fig.S4). Interestingly, Ile (I) and Val (V), which are hydrophobic amino acids, were shown to completely prevent the binding. Ultimately, six point mutations were chosen for experimental testing: I54A, Q66D, V68Y, M115A, Y123D, and R125E, with I54A, Q66D, and M115A expected to be central to the binding epitope. V68Y and Y123D were hypothesized to specifically impact parallel and perpendicular binding modes, respectively, while R125E was anticipated to have minimal interference.

These PD-L1 variants were recombinantly produced and their binding affinities were evaluated using native poly(acrylamide) gel electrophoresis and single bead-based cytometry (Fig.4A, S5, S6). The results indicated moderate binding loss for I54A, Q66D, and M115A, and minimal change for R125E, suggesting the binding epitope is located on the GFCC’ face of the IgV-like domain (Fig.4D). Notably, Y123D’s complete loss of binding suggested the perpendicular binding mode compared to V68Y with moderate affinity loss. Comparing the AlphaFold2 scores with the binding affinity data (Fig.A vs. Fig.4A, Fig.S6) showed that while AlphaFold2 effectively captured major affinity differences, it was less effective in detecting subtle variations. The observed reduction in binding affinity by the Q66D mutation also supported the perpendicular mode, though this conclusion was not definitive.

**Figure 4:**
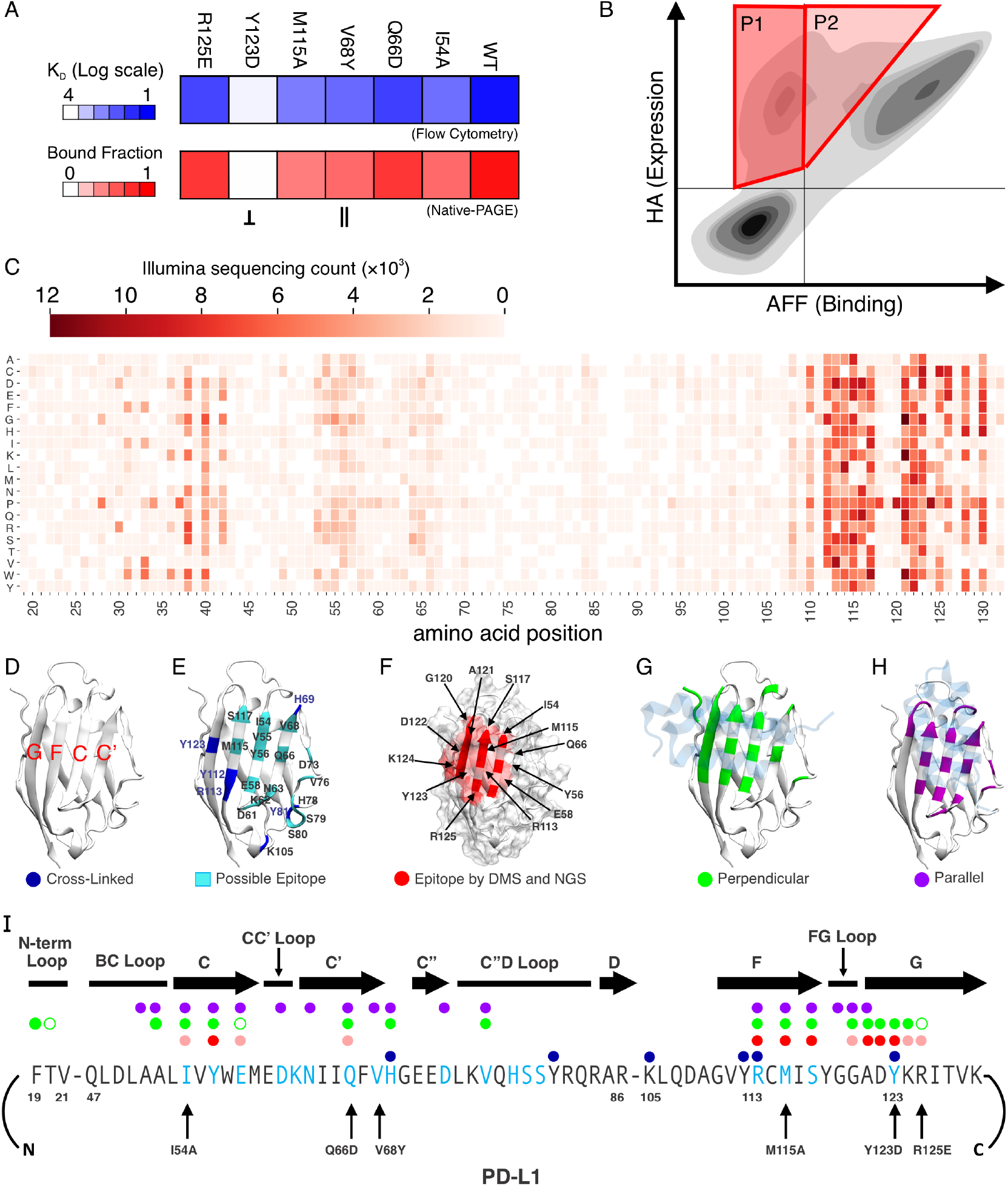
Experimental validation of binding epitope and perpendicular orientation. A) Site-specific PD-L1 mutagenesis and quantification of relative binding activity by flow cytometry and Native-PAGE. B) Cell labelling and sorting for strategy for epitope mapping by deep mutational scanning using a yeast display PD-L1 variant library. C) Heatmap of positions found among PD-L1 variants with reduced/no binding activity. D-G) Epitope mapping analysis results mapped onto the crystal structure of PD-L1. Labels are color coded to match the property analyzed, and indicated along the sequence in I. D), *β*-sheet labels. E) Binding epitope determined by XL-MS analysis (possible epitope in cyan, crosslinked residues in blue). F) Binding epitope determined by DMS and NGS (red). G) Analysis of binding orientation for the perpendicular orientation (green) H) Analysis of binding for the parallel orientation (violet). I) PD-L1 sequence with markings in correponding colors to figures D-H.

Since the site-directed mutagenesis and affinity measurements were not sufficient to differentiate between perpendicular and parallel binding modes, we next turned to crosslinking-coupled mass spectrometry (XL-MS) which identifies the epitope by determining proximity of cross-linkable amino acids side chains within the complex. This involved exposing a sample of the bound PD-L1:Affibody complex to a mass-labeled crosslinker, followed by enzymatic fragmentation and MS/MS analysis to locate cross-linking sites, revealing both the epitope and binding orientation.

The XL-MS analysis revealed ten different proximity-dependent crosslinks between the Affibody and PD-L1, distributed across six PD-L1 amino acids: H69, Y81, K105, Y112, R113, and Y123. These residues form a triangular shape on the GFCC’ face of the IgV-like domain (Fig.4E; Blue). The amino acids within this triangle, including I54, Y56, E58, D61, and others, were therefore implicated as being part of the binding epitope by XL-MS, however again XL-MS alone was not capable of completely excluding either parallel or perpendicular modes.

To provide even higher resolution for experimental epitope mapping, we next turned to deep mutational scanning (DMS) with next-generation sequencing (NGS) by yeast display. This offered a high-throughput approach to epitope mapping and involved displaying a mutated PD-L1 variant library on the surface of yeast cells, and quantifying both PD-L1 variant expression level and Affibody binding strength using fluorescent activated single cell sorting (FACS)/flow cytometry coupled with high-throughput DNA sequencing. The PD-L1 variant library contained all possible single amino acid substitutions within the PD-L1 sequence constructed as a scanning one codon-one amino acid library (Fig. S7, S8). The FACS gating strategy is shown in (Fig.4B). Sorting and deep-sequencing yeasts displaying PD-L1 variants that were highly expressed but which exhibited reduced or no Affibody binding activity (Fig.4B, regions P1 and P2) allowed us to identify the binding epitope with high resolution. The Illumina sequencing read counts of sorted PD-L1 variants were visualized as a heatmap (Fig.4C) and included several surface-exposed residues along with others within the *β*-sandwich of the IgV-like domain, potentially destabilizing PD-L1’s structure (Fig. S9, S10). The binding epitope identified by DMS covering all possible single mutations, showed higher precision compared to the other methods (Fig.4F).

When we compared the computationally suggested perpendicular and parallel modes with the DMS data, both on the primary protein sequence of PD-L1 and its crystal structure with PD-1^52^ (Fig.4G-H), the implicated residue positions validated the perpendicular binding mode, characterized by more significant involvement of the G *β*-strand and less of the BC loop, CC’ loop, C *β*-strand, and C’ *β*-strand. Our findings for the experimental validation of binding epitope and orientation are summarized in Figure 4I.

## Methods

We employed AlphaFold2 Multimer ^23,24^ to predict the structure of the Affibody:PD-L1 complex. To investigate the most probable conformation from the AlphaFold2 predictions, we prepared molecular dynamics (MD) simulations using QwikMD^43^ and carried them out using the GPU-resident version of NAMD 3.0. ^44^ Metrics including RMSD, MM-PBSA, and Dynamic Network Analysis ^45^ were applied to evaluate the stability of the Affibody:PD-L1 complex structure.

To further confirm the correct conformation, we once more employed AlphaFold2 Multimer for a *in silico* mutational scanning. For native PAGE, bead-based cytometry and XL-MS analysis, we produced Affibody and PD-L1 variants in *E. coli*. For DMS, a scanning NNK codon library covering the entire length of PD-L1 was displayed in EBY100 yeast through an Aga2p anchor system. NGS was carried out using paired end 300 bp reads on an Illumina NextSeq2000.

A comprehensive description of methodologies, data availability statements, relevant accession codes, and references can be found in the supporting information of the paper.

## Discussion

Alternative binding scaffolds such as Affibodies are being explored for anti-(PD-L1) therapies. In order to elicit an effective immune response, Affibody binding must target the correct epitope on PD-L1, therefore correctly identifying binding epitopes and providing high resolution methods to compare subtle differences in modes of binding for different binders is crucial. Computational methods offer significant potential in this regard, but frequently they provide ambiguous results.

Here we introduced a methodology combining computational techniques and experimental validation to predict and analyze the structure and interactions of a PD-L1:Affibody complex. Through an integrated approach utilizing AlphaFold2, network analysis of molecular dynamics simulations, and deep mutational scanning we were able to understand and validate the critical interaction regions of a high affinity PD-L1:Affibody complex.

Our findings indicated that AlphaFold2 Multimer, ClusPro, and ZDock yield analogous results, and suggested two potential binding modes referred to as parallel and perpendicular (Fig. 2B,C). We subjected the structures of both modes to MD simulations and used Dynamic Network Analysis of the trajectories to discover that the perpendicular mode exhibited higher stability (Fig. 2F, Fig S2). Notably, traditional MD trajectory analyses such as RMSD and MM-PBSA struggled to distinguish the binding differences between the two modes (Fig. 2D,E). The advantage of the network analysis lies in its foundation on the generalized correlation of atomic motions, making it more adept than standard chemical descriptors in predicting interface interactions.^53^ While conducting *in silico* targeted mutations for epitope mapping, we observed that AlphaFold2 adeptly captured experimentally measured trends in affinity of mutants versus the WT, but failed with small affinity variations (Fig. A, Fig. S6). These results highlight the important role MD simulations can play in successfully refining the AlphaFold2 predictions and elevating the perpendicular binding mode as the most probable predicted mode. Further experimental validation by high resolution cross-linking mass spectrometry and deep mutational scanning confirmed the perpendicular mode as the accurate binding conformation (Fig. 4). Notably, using just AlphaFold2 scoring, less favorable structural conformations, such as the parallel binding mode, still garnered relatively high prediction scores.

In summary, our approach demonstrates the efficacy of combining AlphaFold2 with MD simulations and Dynamic Network Analysis in characterizing binding interfaces. The PD-L1:Affibody complex serves as an illustrative case study, especially considering the multitude of binding regions identified for PD-L1 binders (Fig. 2A). Our computational approach proved robust and was corroborated by an array of experimental techniques. The methodology holds promise for exploring other bimolecular interactions, as it can be easily implemented for other binding interfaces. Moreover, within cancer therapeutics, our technique marks a significant step towards correct epitope targeting for the design enhanced cancer treatments.

## Acknowledgements

Acknowledgments RCB and DEBG are supported by the National Science Foundation under Grant MCB-2143787, and the National Institute of General Medical Sciences (NIGMS) of NIH through the grant R24-GM145965. This work was supported by a grant from the Swiss State Secretariat for Education, Research and Innovation (SERI) to MN.

