## Supplementary Material for "Integrating Dynamic Network Analysis with AI for Enhanced Epitope Prediction in PD-L1:Affibody Interactions"

### Methods

#### PD-L1:Affibody AlphaFold models

The aminoa cid sequences for the mature domain of PD-L1 (UniProt Q9NZQ7, 18-234) and the Affibody (1-60) were used as a single FASTA input. The models for the PD-L1:Affibody complex were predicted using AlphaFold-Multimer[23,24] version 2.3.2, using all 5 available v3 multimer parameter sets, to generate 5 models each, resulting in a total of 25 predictions per sequence pair. The QwikFold VMD's[50] plugin was used to set the experiments and post-process the results, and calculations were run using the Cybershuttle[54] Research Environment deployed at the SDSC Expanse[55] supercomputer. QwikFold was used to align the models for visual inspection, addressing per-residue confidence as measured by pLDDT.[56] The predicted aligned error (PAE) matrices were additionally inspected to assess the confidence in the relative position and orientation of the two major binding conformations: perpendicular or parallel.

##### Using AlphaFold for epitope mapping

For *in silico* mutagenesis aimed at epitope mapping the PD-L1:Affibody complex by mutating the PD-L1 sequence, the selected mutations I54A, V68Y, M115A, Y123D, R125E were used following the same procedure as above. Position Q66 was further scanned for all the twenty common amino acids. The iptm+ptm score for the best ranking model for the perpendicular orientation was plotted as bar graph (Fig A-B)

#### Molecular Dynamics Simulations

The perpendicular and parallel AlphaFold2 predicted models for the PD-L1:Affibody were subjected to refinement and conformational sampling by molecular dynamics simulations, with PD-L1 trimmed to it's IgV domain (18-115). The system was then solvated TIP3P water and neutralized using Sodium atoms as counter-ions, which were randomly arranged in the solvent. Total system size was approximately 40k atoms. The MD simulations in the present study were performed employing the GPU-accelerated molecular dynamics package NAMD.[44] The CHARMM36 force field[57,58] along with the TIP3 water model[59] was used to describe all systems. The simulations were carried out assuming periodic boundary conditions in the NpT ensemble with temperature maintained at 300 K using Langevin dynamics for pressure, kept at 1 bar, and temperature coupling. A distance cut-off of 12.0Å was applied to short-range, non-bonded interactions, whereas long-range electrostatic interactions were treated using the particle-mesh Ewald (PME)[60] method. The equations of motion were integrated using the r-RESPA multiple time step scheme[61] to update the van der Waals interactions every step and electrostatic interactions every two steps. The time step of integration was chosen to be 2 fs for all simulations performed. The first two nanoseconds of the simulations served to equilibrate systems before the 16 production runs for 100 ns.

#### Molecular Dynamics Simulations Analysis

All analyses of MD trajectories were carried out employing VMD,[50] its plugins and TCL scripts, unless stated differently. Analysis outputs were post-processed to generate graphs using Python3[62] libraries, including Matplotlib,[63] Pandas,[64,65] and Seaborn,[66] unless stated differently. Figure panels were assembled with CorelDraw Graphics Suite 2021.

#### Binding Affinity Prediction

MM-PBSA is a computationally efficient method for estimating the binding free energy (ΔGbind) of protein-protein complexes.[67–69] Here, to characterize PD-L1:Affibody coupling, we performed the effective free energy (ΔGeff, neglecting the configurational entropic contribution) binding affinity prediction by MM-PBSA using the CaFE plugin.[70] Molecular Mechanics (MM) was computed with NAMD3,[44] Solvent accessible area with VMD,[50] and Poisson-Boltzmann term was computed with APBS.[71]

#### Dynamic Network Analysis

The dynamical network analysis python package[45] was used to extract correlations of motion from PD-L1:Affibody simulations. A network was defined as a set of nodes, all C$\alpha$, with connecting edges. Edges connect pairs of nodes if corresponding monomers are in contact, and 2 nonconsecutive monomers are said to be in contact if they fulfill a proximity criterion, namely any heavy atoms (non-hydrogen) from the 2 monomers are within 4.5Å of each other for at least 75% of the frames analyzed. A network was defined as a set of C$\alpha$ atoms, linked by edges. Edges connect pairs of nodes if the corresponding monomers are in contact. A contact relies on a proximity criterion: heavy atoms from the two monomers must be within a distance of 4.5Å from each other for at least 75% of the frames analyzed. The dynamical networks were constructed from 20 ns windows of the total trajectories sampled every 200 ps.

#### Cluster analysis

Clustering analysis was performed using the unsupervised k-means algorithm as implemented in CPPTRAJ,[72] with the parameters "randompoint maxit" set to 500 and "sieving" set to 10. The distance metric utilized for clustering was the RMSD of the complex C, N, O, C$\alpha$ and C$\beta$ atoms, and the clustering process concluded when the number of clusters reached 5. The analysis was applied to the last quarter of each of the 16 replica trajectories for each orientation (perpendicular and parallel), after fitting to the complex backbone, excluding the 5 residues from both Affibody's termini.

##### Native contacts for PDB structures

To reveal insights into the PD-L1:Affibody complex assembly we performed an analysis of PD-L1 (UniProt Q9NZQ7) structures found in the Protein Data Bank (PDB), with interactions with other proteins. The native contacts identified after superposition of available structures, with contacts defined by amino acid residues within a 4Å radius of the PD-L1. Residues with contacts are highlighted in Fig.2. The structures considered were: 3BIK, 3FN3, 3SBW, 4Z18, 4ZQK, 5GGT, 5GRJ, 5IUS, 5J89, 5J8O, 5JDR, 5JDS, 5N2D, 5N2F, 5NIU, 5O45, 5O4Y, 5X8L, 5X8M, 5XXY, 6NM7, 6NM8, 6NNV, 6NOJ, 6NOS, 6PV9, 6R3K, 6RPG, 6VQN, 6YCR, 7BEA, 7C88, 7CZD, 7DY7, 7NLD, 7OUN, 7SJQ, 7TPS.

#### Plasmids available on addgene:

Addgene plasmid #157674: pET28a-ybbR-His-ELP(MV7E2)3-FLN-SpyCatcher.

#### Cloning of PD-L1-HIS and PD-L1-HIS-SpyTag

The DNA sequence of the extracellular domain of human PD-L1 was chemically synthesized with optimal codons for production in *E. coli* (GeneArt, Thermo Fisher Scientific) and introduced into a pET28a vector via NdeI and XhoI restriction sites to generating a new vector pET28a-PD-L1-HIS. The plasmid contents were confirmed by further Sanger DNA sequencing analysis. A SpyTag was further introduced at the C-terminus of PD-L1 by PCR using primers #1 and #2 (Table S2) based on the plasmid pET28a-PD-L1-HIS and following Gibson assembly with master mix (NEB) generating a new vector pET28a-PD-L1-ECD-HIS-SpyTag, which was confirmed by further DNA sequencing analysis.

#### Cloning of PD-L1-HIS-SpyTag Mutants

For the mutational analysis, six point-mutations (I54A, Q66D, V68Y, M115A, Y123D and R125E) were designed and incorporated by site directed mutagenesis using the

Q5® Site-Directed Mutagenesis kit (NEB) with primers #3 and #4 for **I54A**, #5 and #6 for **Q66D**, #6 and #7 for **V68Y**, #8 and #9 for **M115A**, #10 and #11 for **Y123D**, and #11 and #12 for **R125E** (Table S2), generating new plasmids pET28-PD-L1-SpyTag-**I54A**, **-Q66D**, **-V68Y**, **-M115A**, **-Y123D** and **-R125E**, which was confirmed by further DNA sequencing analysis.

#### Expression, Refolding, and Purification of PD-L1 Variants

The plasmid with the sequence of the PD-L1 variant was introduced into competent *E. coli* BL21(DE3) strain. Recombinant cells were cultured in 5 ml of Luria-Bertani (LB) medium with 50 $\mu$g ml$^{-1}$ kanamycin at 37 °C overnight. The culture was transferred to 50 mL of Terrific broth (TB) medium with 50 $\mu$g ml$^{-1}$ kanamycin and cultivated at 37 °C and 200 rpm until an optical density at 600 nm (OD600) of 0.8-1.0 was reached. The expression of recombinant protein was induced by the addition of 1.0 mM isopropyl-$\beta$-D-thio-galactopyranoside (IPTG) and the culture was further incubated at 37 °C and 200 rpm for 9 hrs. The cells were harvested by centrifugation at 4,000 g for 20 min at 4 °C. The harvested cell pellet was resuspended in a denaturing lysis buffer (10 mM Tris-Cl, and 8M urea; pH 8). Resuspended cells were placed on ice and disrupted for 15 min using a sonic dismembrator using a 3 s on: 5 s off pattern to allow cooling between each pulse. The lysate was centrifuged at 14,000 g for 20 min at 4 °C. The supernatant was collected and incubated with Ni-NTA resin for 30 min at room temperature to allow the His6-tagged proteins to bind to the Ni-NTA resin. Then, the mixture was loaded onto a column. The resin was washed with 10–20 resin volumes of wash buffer (20 mM imidazole, 10 mM Tris-Cl, and 8M urea; pH 8). Recombinant proteins were eluted in the elution buffer (500 mM imidazole, 10 mM Tris-Cl, and 8M urea; pH 8). The eluted protein solution was serially dialyzed to 8 M, 4 M, 2 M, and 0 M Urea with 5% glycerol, 5% sucrose, 1% arginine, 0.5 mM NaCl in 20 mM Tris-Cl (pH 7.4), and finally to 1x PBS buffer. Precipitation during dialysis was removed by centrifugation at 14,000 g for 20 min at 4 °C and supernatant was further purified by SEC column.

#### Cloning of AFF-HIS and ybbR-HIS-ELP-FLN-Anti-PD-L1-AFF (L-AFF)

DNA sequence of Anti-PD-L1 Affibody (AFF) was chemically synthesized based on the codon usage of E. coli (GeneArt, Thermo Fisher Scientific) and introduced into pET28a vector via NdeI and XhoI restriction sites generating a new vector pET28a-Anti-PD-L1-AFF-HIS (for preparation of AFF-HIS), which was confirmed by further DNA sequencing analysis. To immobilize AFF on the PS beads for flow cytometry analysis, ybbR tag and linker was introduced at the N-terminus of AFF (for preparation of L-AFF) by PCR using primers #13 and #14 based on the plasmid pET28a-Anti-PD-L1-AFF-HIS and using primers #15 and #16 based on the plasmid #157674 (Addgene) (Table S2). Two PCR products were assembled into a new vector pET28a-ybbR-HIS-ELP-FLN-Anti-PD-L1-AFF by Gibson assembly with master mix (NEB), which was confirmed by further DNA sequencing analysis.

#### Expression and Purification of AFF-His and L-AFF

The plasmid with the sequence of AFF-His or L-AFF was introduced into competent E. coli BL21(DE3) strain. Recombinant cells were cultured in 5 ml of Luria-Bertani (LB) medium with 50 $\mu$g ml$^{-1}$ kanamycin at 37 °C overnight. The culture was transferred to 50 mL of Terrific broth (TB) medium with 50 $\mu$g ml$^{-1}$ kanamycin and cultivated at 37 °C and 200 rpm until an optical density at 600 nm (OD600) of 0.8-1.0 was reached. The expression of recombinant protein was induced by the addition of 0.5 mM IPTG and the culture was further incubated at 20 °C and 200 rpm for 9 hrs. The cells were harvested by centrifugation at 4,000 g for 20 min at 4 °C. The cells were harvested by centrifugation at 4,000 g for 20 min at 4 °C. The harvested cell pellet was resuspended in lysis buffer (50 mM Tris, 50 mM NaCl, 0.1% Triton X-100, 5 mM MgCl2; pH 8.0), and disrupted with a sonic dismembrator. The lysate was centrifuged at 14,000 g for 20 min at 4 °C. The supernatant was collected and incubated with Ni-NTA resin, loaded onto a column, washed with wash buffer (1x PBS with 20 mM imidazole; pH 7.4), and eluted in elution buffer (1x PBS with 500 mM imidazole; pH 7.4). The eluted protein solution was further purified by SEC column.

#### Native-PAGE Analysis

Binding behavior between AFF and PD-L1 mutants (WT, I54A, Q66D, V68Y, M115A, Y123D and R125E) were screened by Native-PAGE. 5 $\mu$L of 10 $\mu$M L-AFF was mixed with 5 $\mu$L of each 10 $\mu$M PD-L1 mutants, incubated several hours at RT, and then total solution was run in Native-PAGE. Protein bands were visualized by Coomassie staining. Bound and unbound fraction of PD-L1 were calculated based on the intensity of stained protein bands.

#### Flow Cytometry Analysis

The binding affinity between Anti-(PD-L1)-AFF and PD-L1 mutants was analyzed using the Attune NxT (Thermo Fisher Scientific) flow cytometer equipped with a 488 nm and a 561 nm laser. L-AFF was immobilized onto the surface of amine-functionalized PS beads via ybbR Tag. The amine groups reacted to a NHS group from sulfosuccinimidyl 4-(N-maleimidomethyl)cyclohexane-1-carboxylate (sulfo-SMCC; Thermo Fischer Scientific) in 50 mM HEPES buffer pH 7.5 for 30 min. The thiol group from Coenzyme A (CoA, 200 $\mu$M) reacted to a maleimide group from sulfo-SMCC in coupling buffer (50 mM sodium phosphate, 50 mM NaCl, 10 mM EDTA, pH 7.2) for 2 hrs. Finally, the ybbR-tagged protein L-AFF was immobilized onto the surface using SFP-mediated ligation to CoA in Mg2+ supplemented 1x PBS buffer. This resulted in covalent immobilization of AFF to PS beads. Protein-immobilized beads were extensively washed and kept in 1x PBS buffer prior to immediate use. GFP-labeled PD-L1 mutants were prepared by conjugating SpyTag of PD-L1 mutants to GFP-SpyCatcher. L-AFF immobilized beads were incubated in GFP-labeled PD-L1 mutants' solution with different concentrations ranging from 0.008 nM to 625 nM for 1-2 hrs at RT. After washing, shift of fluorescence from GFP was recorded and plotted against the concentration of PD-L1 mutants to derive the dissociation constant between PD-L1 mutants and L-AFF.

#### XL-MS Epitope Mapping

The conformational epitope mapping for PD-L1 and Anti-PD-L1-AFF was performed by chemical cross-linking and high-resolution mass spectrometry (XL-MS; CovalX, Switzerland). 10 $\mu$L of PD-L1 (6 $\mu$M) was mixed with 10 $\mu$L of AFF (6 $\mu$M) to obtain PD-L1/AFF mix with a final concentration of 3 $\mu$M. Then, 2 $\mu$L of deuterated cross-linker disuccinimidyl suberate (DSS d0/d12, 2 mg/mL in DMF) was added to the protein mixture and the solution was incubated for 180 min at RT to complete the cross-linking reaction. After cross-linking the reaction was stopped with 20 mM ammonium bicarbonate, the samples were submitted to reduction, alkylation, and proteolysis with five different enzymes (trypsin, chymotrypsin, ASP-N, elastase, and thermolysin). After enrichment of the cross-linked peptides, the samples were analyzed by high-resolution mass spectrometry (nLC-Q-Exactive Orbitrap MS). The NHS groups of DSS reacts only with positively charged amino groups or hydroxyl groups including Arg, His, Lys, Ser, Thr, and Tyr. Specific amino acid residues that were cross-linked were identified by tandem MS/MS analysis. Based on these cross-linked amino acids and the crystal structure of PD-L1, possible binding epitope was proposed.

#### Cloning for Yeast Surface Display of PD-L1

For the mutational analysis, PD-L1 sequence was amplified by PCR using primers #17 and #18 (Table S2) based on the plasmid pET28a-PD-L1-ECD-HIS and introduced via NheI and BamHI restriction site into the yeast plasmid pYD1 for surface display (Addgene plasmid #73447) generating a new vector pCHA-HA-PD-L1-ECD-HIS-Xpress-Aga2p, which was confirmed by further DNA sequencing analysis.

#### Site-Saturation PD-L1 Libraries

Sequence of PD-L1-ECD was divided into two regions; IgV-like domain (F19 to A132; 114 aa) and IgC-like domain (P133 to R238; 106 aa). Site-saturation mutagenesis libraries spanning all 114 positions for IgV-like domain (Library A) and for all 106 positions for IgC-like domain (Library B) encoding all the 19 possible amino acid mutations were produced by Twist Bioscience. The backbone template was prepared by digestion of plasmid pCHA-HA-PD-L1-ECD-HIS-Xpress-Aga2p with NheI and BamHI. After agarose gel electrophoresis, purified backbone template was mixed with each of synthesized PD-L1 Library A and B with a ratio of 1:10. Then, the mixture was directly transformed into Saccharomyces cerevisiae EBY100 following a typical lithium acetate transformation procedure for introducing site=saturation mutagenesis library into backbone template via endogenous homologous recombination. Right after transformation, serial dilutions were plated on synthetic defined (SD) agar 2% (w/v) glucose plates lacking tryptophan (-Trp) to count the number of transformants. Remaining transformation reaction solution was further grown in liquid SD glucose -Trp medium for 48 h at 30 °C with continuous shaking at 200 rpm. Then, cultured yeast library was harvested, prepared as 50% glycerol stock and stored at -80 °C for further analysis.

#### Yeast Surface Display

Saccharomyces cerevisiae EBY100 transformants harboring the plasmid pCHA-HA-PD-L1-ECD-HIS-Xpress-Aga2p or PD-L1 yeast library A and B were cultivated in SD-TRP liquid medium with 2% glucose for 24 h at 30 °C with continuous shaking at 200 rpm. Protein expression and protein display were then induced by transferring the culture to a fresh pH-buffered liquid medium (0.1 M Potassium phosphate pH 7.0) lacking tryptophan containing 0.2% (w/v) glucose and 1.8% (w/v) galactose and by shaking for 24 h at 30 °C.

#### Cell Sorting with Dual Labelling

PD-L1 libraries are displayed with N-terminal HA tag. Therefore, the expression of PD-L1 was labelled via HA-tag and the binding of AFF-HIS was labelled via HIS-tag. Yeast cells displaying the PD-L1 libraries were incubated with 4 nM of AFF-HIS in 1x PBS containing 0.1% BSA for ¿ 1 h at RT with shaking. After washing with 1x PBS containing 0.1% BSA, cells were incubated with a mixture of primary antibodies, HA tag recombinant rabbit monoclonal antibody (RM305; Thermo) and 6x-His tag monoclonal antibody (HIS.H8; Sigma) in 1x PBS containing 0.1% BSA for 1 h at RT. After washing, cells were further incubated with a mixture of secondary antibodies, Goat anti-rabbit IgG (H+L) cross-adsorbed secondary antibody, alexa Fluor™ 488 (Thermo) and Goat anti-Mouse IgG (H+L) highly cross-adsorbed secondary antibody, Alexa Fluor™ 594 (Thermo) in 1x PBS containing 0.1% BSA for 1 h on ice. Finally, dually labelled cells were washed with ice-cold 1x PBS containing 0.1% BSA for further analysis. Cells were sorted by FACSMelody™ Cell Sorter (BD Bioscience). Cells with high expression level with decreased or no binding were sorted, transferred to SD -TRP liquid medium with 2% glucose and 50 $\mu$g ml-1 ampicillin, cultivated for 48 h at 30 °C, harvested, and prepared as 15% glycerol stock and stored at -80 °C for further analysis.

#### Illumina Sequencing and Data Analysis

Plasmids were extracted from the sorted cells using zymolyase (Zymo Research, Irvine, USA) and GeneJET Plasmid Miniprep Kit (Thermo Scientific). Regions of the PD-L1 IgV- and IgC-like domain were amplified by the first PCR step and then indexes and adapters were added for Illumina sequencing by the second PCR step with the primer sets from IDT for Illumina–DNA/RNA UD Indexes Plate A and NEBNext® Ultra™ II Q5® Master Mix (NEB). Final products were purified using AMPure XP beads (Beckman Coulter) and Illumina sequencing was performed with the Illumina NextSeq2000 for paired end 300 bp (PE 2x 300 bp, 600 cycles) (Functional Genomics Center Zurich). The following data analyses were performed at sciCORE (http://scicore.unibas.ch/)

scientific computing center at University of Basel. Briefly, sequencing results from the paired end were combined after trimming reads presenting a quality under 20 and translated into amino acid sequences. Each of identical amino acid sequences only with one mutation (or wild type) were grouped and counted to calculate the ratio in the sorted population (high expression with low/no binding) compared to the initial population.

#### Data availability

All files are available upon request, including AlphaFold run scripts and output models; molecular dynamics initial structures, configuration files, trajectories, and analysis scripts.

Supplementary Table 1: K-means cluster analysis of the 16, 100ns molecular dynamics simulations of the PD-L1:Affibody complex, after fitting to PD-L1's backbone of the initial AlphaFold model.

| Orientation | Cluster | Fraction | AvgDist | Stdev | AvgCDist |
|---|---|---|---|---|---|
| perpendicular | 1 | 0.315 | 1.359 | 0.235 | 1.309 |
| | 2 | 0.239 | 1.405 | 0.264 | 1.200 |
| | 3 | 0.201 | 1.499 | 0.316 | 1.770 |
| | 4 | 0.201 | 1.499 | 0.316 | 1.770 |
| | 5 | 0.201 | 1.499 | 0.316 | 1.770 |
| parallel | 1 | 0.564 | 1.591 | 0.361 | 3.765 |
| | 2 | 0.323 | 1.924 | 0.589 | 3.839 |
| | 3 | 0.050 | 1.714 | 0.521 | 4.554 |
| | 4 | 0.043 | 1.463 | 0.440 | 4.433 |
| | 5 | 0.019 | 2.026 | 0.724 | 5.339 |

Supplementary Table 2: Primers

| Number | Sequence |
|---|---|
| 1 | 5’-GTCATGGTTGATGCATACAAGCCGACGAAGTAACTCGAGTAAGATCCGGCTG-3’ |
| 2 | 5’-GGCTTGTATGCATCAACCATGACAATATGGGCGCTACCGTGATGATGATGGTGATGGCTAC-3’ |
| 3 | 5’-GGCAGCACTGGCTGTTTATTGGG-3’ |
| 4 | 5’-AGATCCAGCTGTTTTTCAAC-3’ |
| 5 | 5’-AAACATCATCGATTTTGTGCACGG-3’ |
| 6 | 5’-TTATCTTCCATTTCCCAATAAAC-3’ |
| 7 | 5’-AAACATCATCCAGTTTTATCACGGTGAAGAG-3’ |
| 8 | 5’-TTATCGTTGTGCTATTAGCTATGGTGGTGCCG-3’ |
| 9 | 5’-ACACCGGCATCCTGCAGT-3’ |
| 10 | 5’-CGATGATAAACGTATTACCGTTAAAGTGAATGC-3’ |
| 11 | 5’-GCACCACCATAGCTAATC-3’ |
| 12 | 5’-CGATTACAAAGAAATTACCGTTAAAGTGAATGC-3’ |
| 13 | 5’-GGGCTCCGGTTCTGGTTCCGTGGACGCCAAATATGCCAAAG-3’ |
| 14 | 5’-GCAGCCGGATCTTACTCGAGTTATTTCGGTGCCTGGCTATCATTC-3’ |
| 15 | 5’-GGAACCAGAACCGGAGCCCGGAGCCGGTTTAACGGTAAC-3’ |
| 16 | 5’-CTCGAGTAAGATCCGGCTGC-3’ |
| 17 | 5’-CAAGCTAGCTTTACCGTGACCGTTCCGAAAG-3’ |
| 18 | 5’-CTTGGATCCGTGATGATGATGGTGATGGCTAC-3’ |

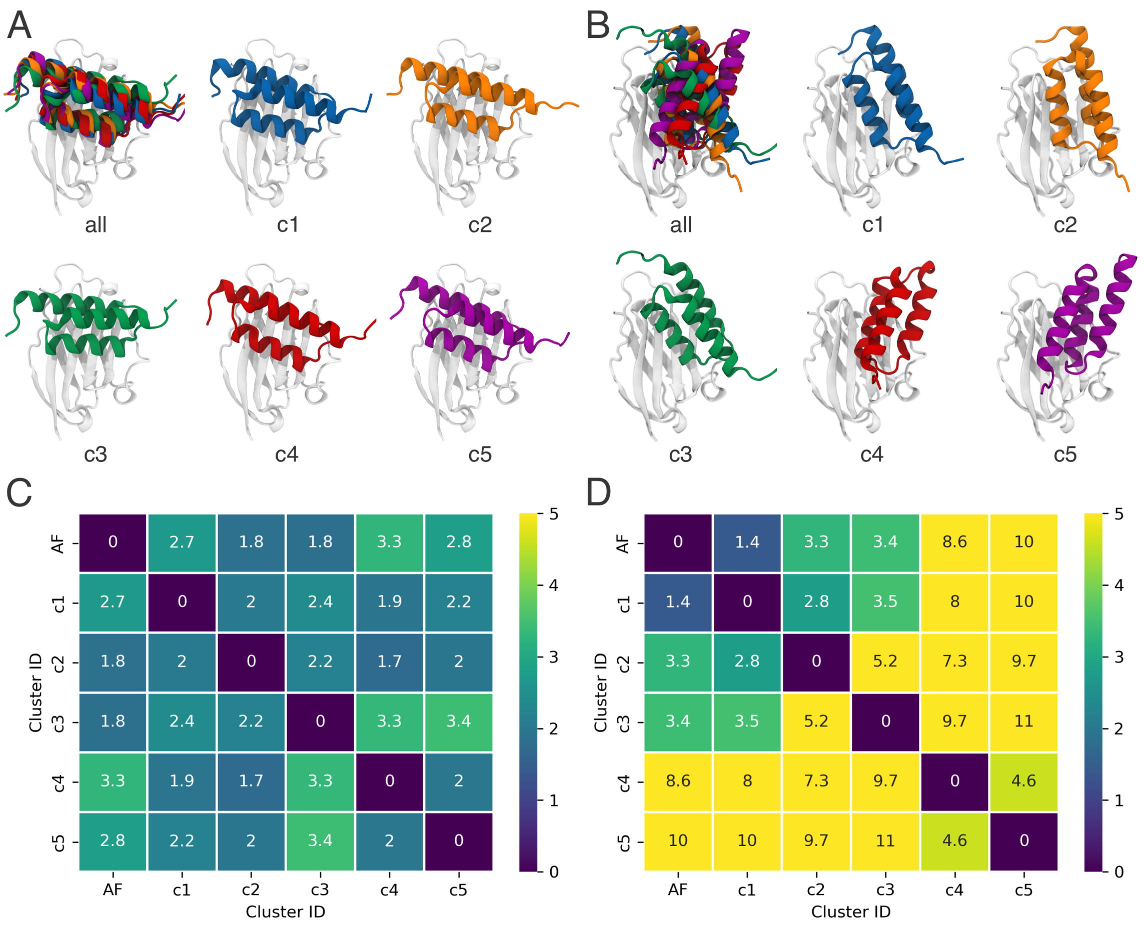


Supplementary Figure S1: K-means cluster analysis of the 16, 100ns molecular dynamics simulations of the PD-L1:Affibody complex, after fitting to PD-L1's backbone of the initial AlphaFold model. A and B refer to the perpendicular and parallel conformations, respectively, with matching colors for each cluster. C and D show the RMSD matrix between the representative structures of each cluster for the perpendicular and parallel conformations, respectively. The Affibody's backbone RMSD to the initial AlphaFold model as well as each cluster's prevalence.

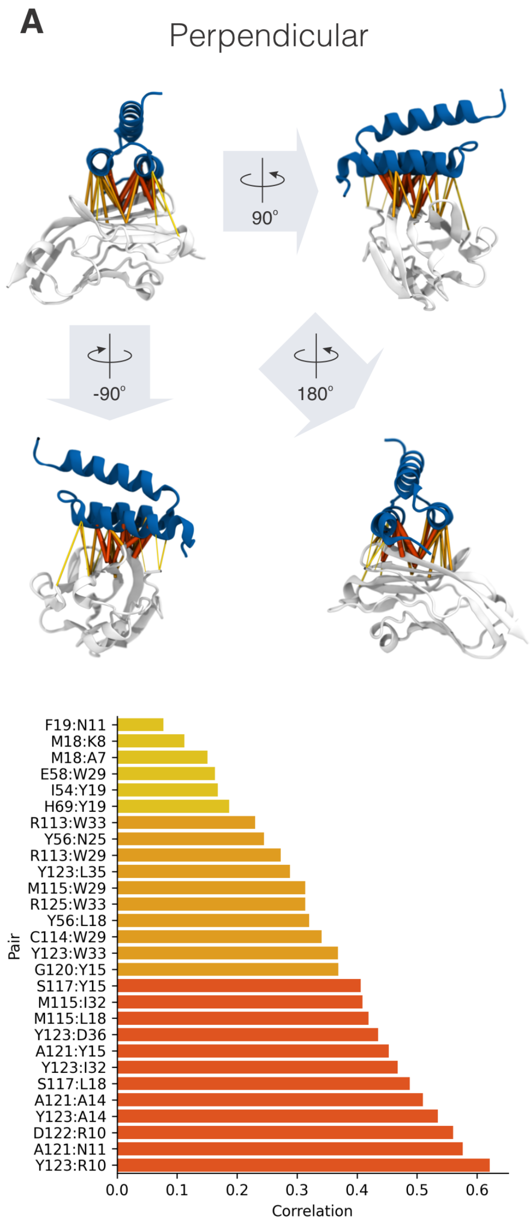


Supplementary Figure S2: Correlations of motion from PD-L1:Affibody simulations from dynamic network analysis.[45] A network was defined as a set of nodes (C$\alpha$) with edges connecting node pairs, for residues in contact for at least 75% of the analyzed frames. At the top, the figure illustrates correlations between residue pairs with lines connecting PD-L1 and Affibody representations depicted as cartoons. The figure showcases various views for both Perpendicular (A) and Parallel (B) orientations. In these representations, the PD-L1 IgV-like domain is depicted in white, while the Affibody is shown in blue. Correlation lines are displayed with varying widths and colors, indicating three categories: low ($< 0.2$), medium ($>= 0.2, < 0.4$), and high($>= 0.4$). These categories align with the residue pair correlation bar plots presented at the bottom of the figure.

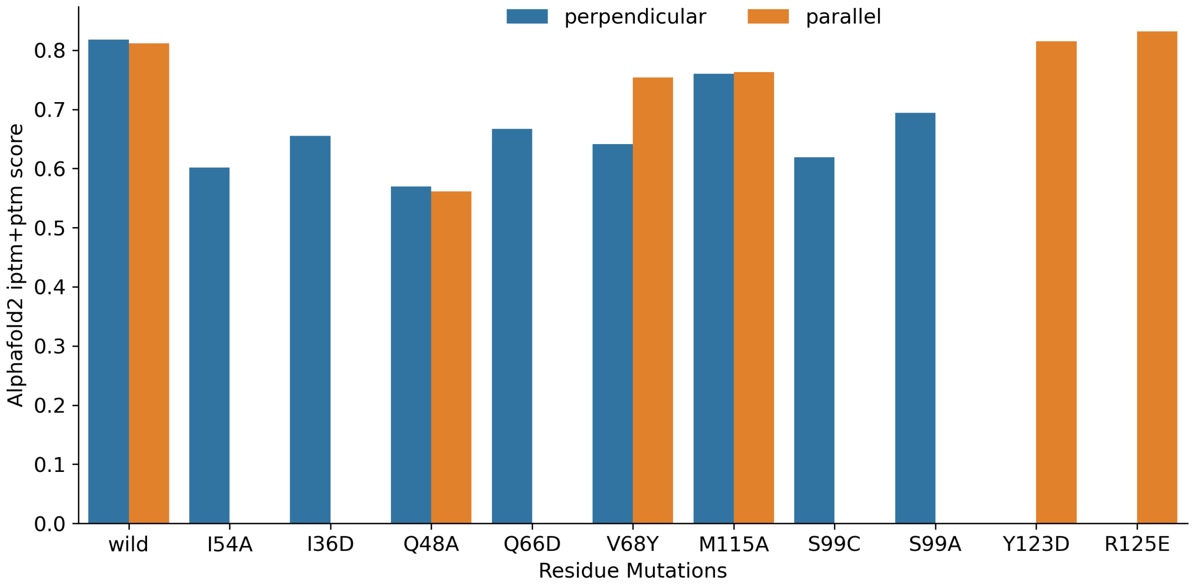


Supplementary Figure S3: Epitope mapping to verify which residues that are key for each of the binding modes. AlphaFold2's iptm+ptm score for the best ranking model for the complex while screening of selected PD-L1 mutations within the predicted perpendicular (blue) and parallel (orange) binding interfaces. The scanned residues were: I54A, Q66, V68Y, M115A, Y123D, and R125E. The absence of a bar indicates no model was generated for that particular orientation.

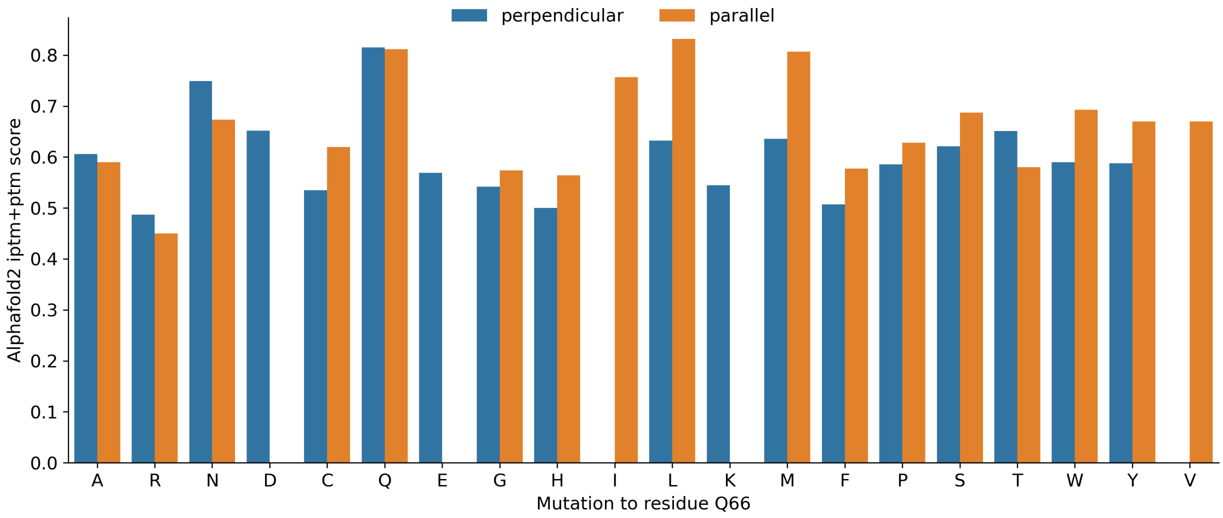


Supplementary Figure S4: Epitope mapping to verify which residues that are key for each of the binding modes while screening PD-L1 mutations for the Q66 position. The figure shows AlphaFold2's iptm+ptm score for the best ranking model for the complex within the predicted perpendicular (blue) and parallel (orange) binding interfaces. The absence of a bar indicates no model was generated for that particular orientation.

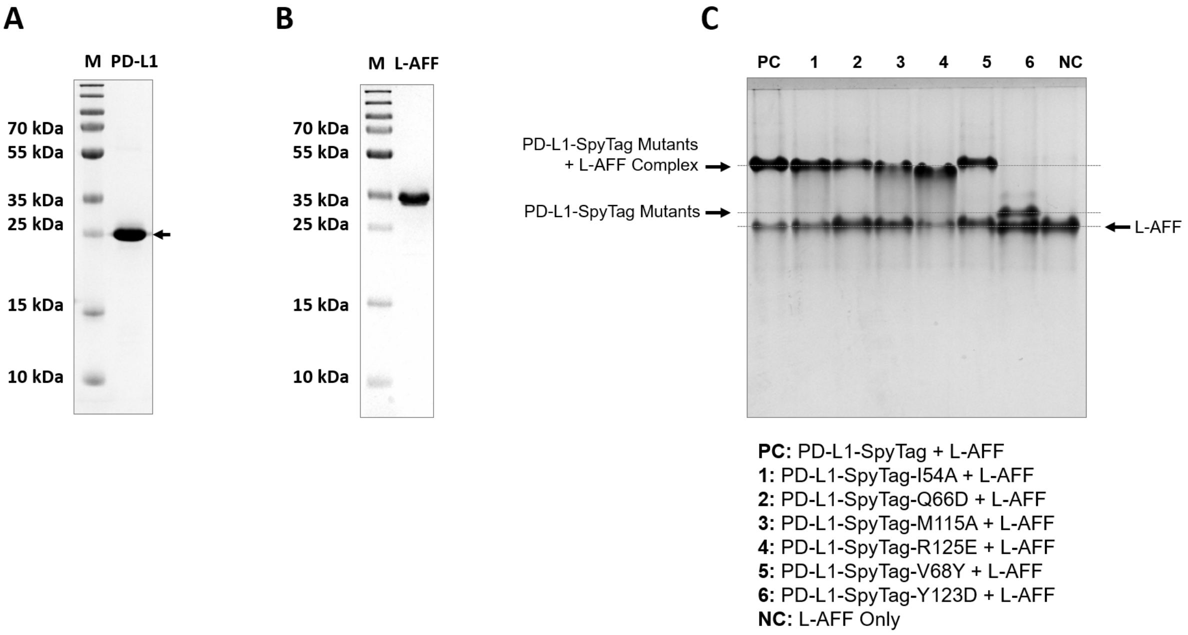


Supplementary Figure S5: Protein preparation and analysis. A) Purified PD-L1 and B) L-AFF analyzed by SDS-PAGE. C) Native PAGE analysis of PD-L1-SpyTag mutants with L-AFF.

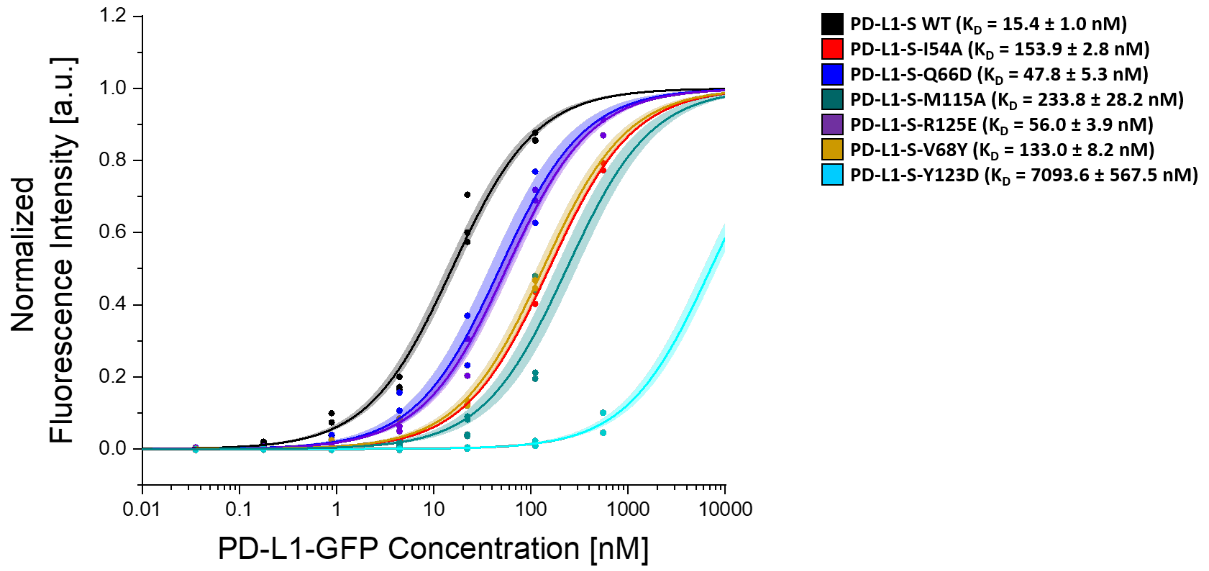


Supplementary Figure S6: Binding affinity analyzed by flow cytometry. AFF-immobilized PS beads were titrated with different concentration of GFP-conjugated PD-L1 mutants. Bound fraction which is labeled as normalized fluorescence intensity was plotted against PD-L1 mutants' concentration.

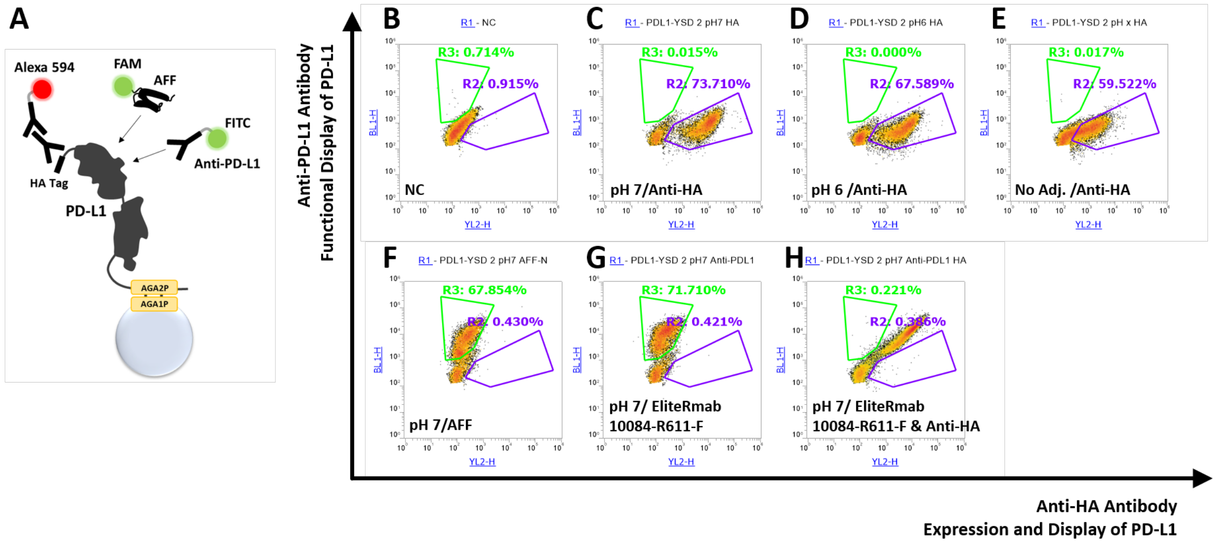


Supplementary Figure S7: Yeast display of PD-L1. A) Schematic illustration of PD-L1 surface display on the yeast cells and the labelling scheme of anti-HA tag with regards to expression/display level of PD-L1 and of anti-PD-L1 AFF and/or monoclonal antibody (CD274 (PD-L1, B7-H1) Monoclonal Antibody (MIH1); Thermo Scientific) with regards to the functional expression/display level of Pd-L1. Flow cytometric cytogram after labelling with anti-HA tag antibody of B) negative control (before expression of PD-L1), C) PD-L1 expression with buffered medium at pH 7, D) with buffered medium at pH 6, E) with non-buffered medium. F) Flow cytometric cytogram after labelling with anti-PD-L1 AFF. G) Flow cytometric cytogram after labelling with anti-PD-L1 monoclonal antibody. H) Dual labelling with anti-HA tag antibody and anti-PD-L1 monoclonal antibody.

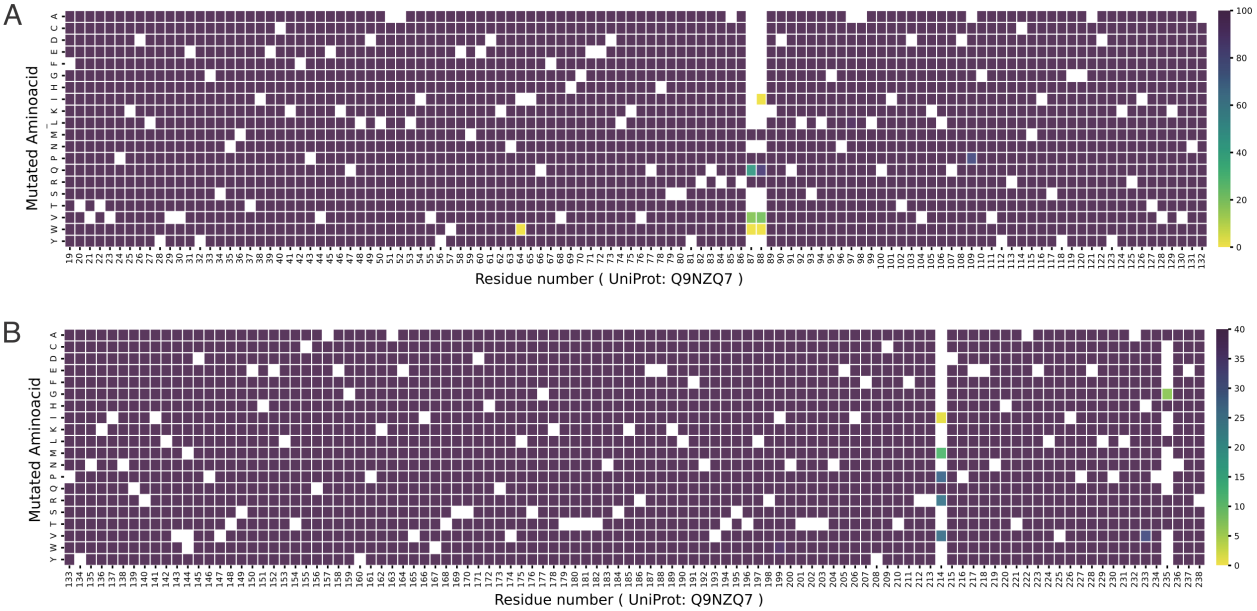


Supplementary Figure S8: Quality control of PD-L1 Library with Illumina sequencing. A) PD-L1 Library A from the total read numbers of 7.2 M with a coverage of 98.5% (2245/2280, ¿100 counts) containing 23.1% of WT. B) PD-L1 Library B from the total read numbers of 2.4 M with a coverage of 98.4% (2086/2120, ¿40 counts) containing 43.1% of WT.

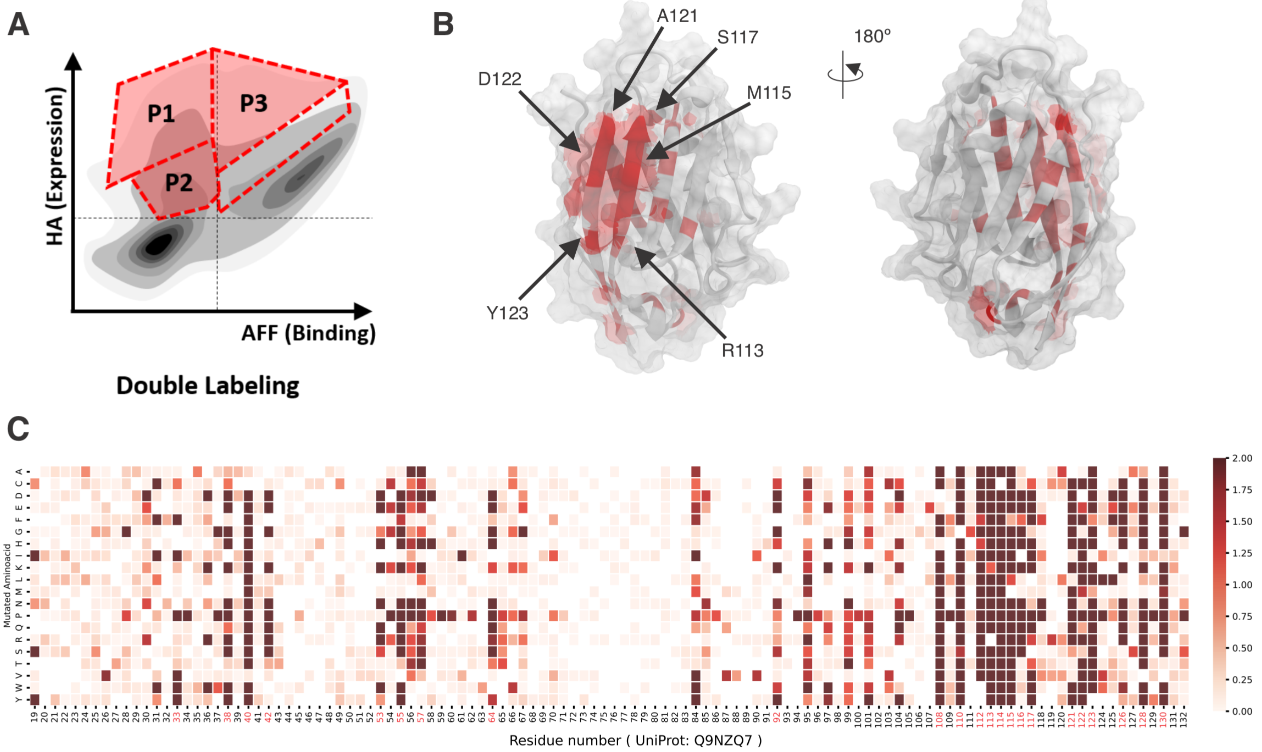


Supplementary Figure S9: Epitope mapping via DMS & Illumina sequencing with dual labelling using PD-L1 Library A using different gates. A) Cells were collected from P1, P2, and P3 fractions. B) Selected amino acid positions were indicated on the crystal structure of PD-L1 IgV-like domain (red). C) Illumina sequencing results presented by heat map. Mutation point with high value of relative probability were selected (residue numbers in red).

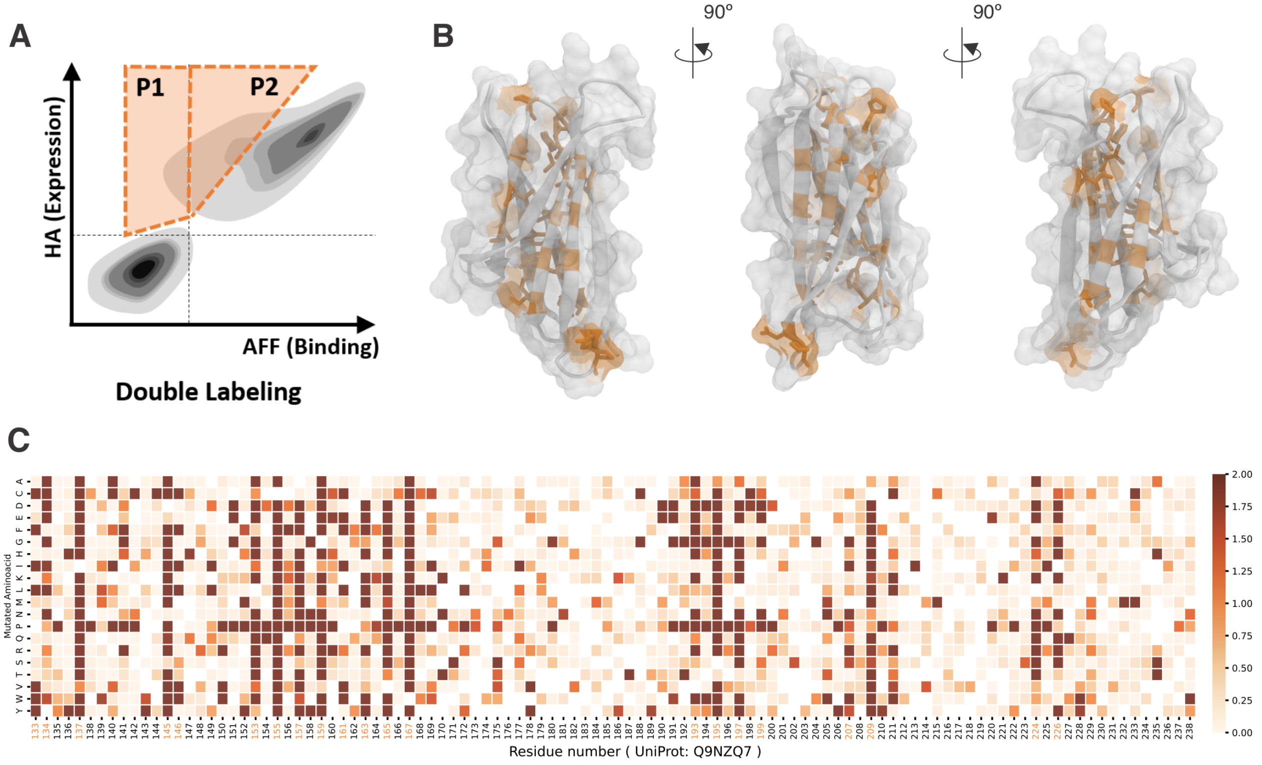


Supplementary Figure S10: Epitope mapping via DMS & Illumina sequencing with dual labelling using PD-L1 Library B. A) Cells were collected from P1 and P2 fractions. C) Selected amino acid positions were indicated on the crystal structure of PD-L1 IgC-like domain (orange). B) Illumina sequencing results presented by heat map. Mutation point with high value of relative probability were selected (residue numbers in orange)
